# Effects of active dry yeast on growth performance, rumen fermentation characteristics and slaughter performance in beef cattle

**DOI:** 10.1101/784082

**Authors:** Siqiang Liu, Mei Yuan, Kun Kang, Zhisheng Wang, Lizhi Wang, Bai Xue, Gang Tian, Huawei Zou, Ali Mujtaba Shah, Xiangfei Zhang, Peiqiang Yu, Hongze Wang, Quanhui Peng

**Author notes:** These authors contributed equally to this work (Siqiang Liu, Mei Yuan). **Notes:** The authors declare no competing financial interest.

## Abstract

This study aimed to investigate the effects of active dry yeast (ADY) on rumen microbial composition and slaughter performance of beef cattle. Thirty-two finishing beef cattle (simmental crossbred cattle ♂ × cattle-yaks ♀), with an average body weight of 110 ± 12.85 kg, were randomly assigned to one of four treatments: the low plane of nutrition group (Control), low plane of nutrition group + ADY 2 g/head/d (ADY2), low plane of nutrition group + ADY 4 g/head/d (ADY4) and high plane of nutrition group (HPN). ADY supplementation increased average daily gain (*P*<0.001), and the carcass weight of ADY4 group had no significant difference with HPN group (*P*>0.05). The serum glutamic-pyruvic transaminase activity in control and ADY4 group was higher than HPN group (*P*=0.001). The neutral detergent fiber (*P*=0.022) and acid detergent fiber (*P*=0.043) digestibility in HPN group was greater than control, but no difference was obtained among ADY2, ADY4 and HPN group (*P*>0.05). The rumen ammonium nitrogen content in control was greater than ADY2 and ADY4 group (*P*=0.003), and no difference was obtained ADY2, ADY4 and HPN group (*P*>0.05). The propionic acid content in the rumen in ADY2, ADY4, and HPN group were greater than control group (*P*<0.001). The simpson (*P*=0.014) and shannon (*P*=0.045) indexes in control and HPN group were greater than ADY4 group. At the phylum level, the relative abundance of Firmicutes in the HPN group was higher than ADY4 group (*P*=0.015). At the genus level, HPN and ADY4 were clustered together, and the relative abundance of *Ruminococcaceae UCG-002* in ADY4 group was higher than control and HPN group (*P*=0.004). In conclusion, supplementation ADY 4 g/head/d shift the rumen microbial composition of beef cattle fed low plane of nutrition to a more similar level with cattle fed with HPN diet, produced comparable carcass weight with HPN diet.

## Introduction

Animal husbandry is now a dynamic and highly developed industry, which is mainly driven by population growth, income growth and urbanization[1]. The rapid development of the animal husbandry is due to the increased demand for livestock products such as meat, milk and eggs, which makes the current production system face challenges. Animal production directly or indirectly contributes 9% of total CO_2_ emissions, 37% of methane emissions and 65% of nitrous oxide emissions, resulting in global warming[2]. China imports more than 90 million tons of soybean every year. Environmental pollution and shortage of protein feed resources are the main factors that restrict the sustainable development of animal husbandry in China.

Beef cattle is an important source of high-quality protein and human economy. As the global population increasing, the competition between people and livestock like land, water and food is becoming increasingly fierce, especially in beef cattle breeding industry[3]. The elevated feed efficiency in cattle could reduce feed consumption while maintaining higher or equal production performance. In addition, cattle with high feed efficiency can not only reduce methane emissions, but also reduce fecal excretion[4, 5]. Therefore, improving the feed efficiency can also reduce the negative impact on the environment.

In recent years, with the increase of people’s attention to animal product safety, quality and environmental issues, more and more green feed additives have been used in livestock breeding, and the effect is obvious. Active dry yeast (ADY) is used in ruminant as a feed additive to prevent health disorders and to improve performance [6]. It is suggested the role of ADY in ruminant diets was to decrease the risk of low ruminal pH[7, 8] and to have fewer liver abscesses. Jiao et al. reported that dietary supplementation of 1.5 g/d ADY (1.71×10^10^ CFU/g) promoted the digestibility of nutrients in small intestine and whole-digestive tract of beef cattle fed high-grain diet, but had no significant effect on growth performance and carcass quality was obtained[9]. In addition, adding 1.5 g/d ADY (1.70×10^10^ CFU/g) to the diet had no effect on the growth performance of fattening beef cattle, but reduced the rumen acidosis incidence and liver abscess[10, 11].

ADY can be used as a feed additive for ruminants to optimize rumen fermentation characteristics hence prevent health problems, however there is little study on the effects of ADY on rumen microbial composition and slaughter performance in beef cattle. We hypothesized that the addition of ADY will change rumen microbial composition and promote beef cattle slaughter performance to a comparable slaughter performance with beef cattle fed high plane of nutrition diet. Therefore, the objectives of this experiment were to determine the effects of supplementation of ADY to low plane of nutrition diet on the growth performance, rumen microbial composition and slaughter performance of beef cattle.

## Materials and methods

All procedures involving animal care were under the approval of the Sichuan Agricultural University Institutional Animal Care and Use Committee.

### Experimental Design, Animals, and Dietary Treatments

Thirty-two fattening beef cattle (simmental crossbred cattle ♂ × cattle-yaks ♀), mean body weight (BW) ± standard deviation: 110 ± 12.85 kg, were randomly assigned to one of four groups with 8 replicates per group and one beef cattle per replicate. Three groups were designed as low plane of nutrition (Control) and one group was regarded as a high plane of nutrition (HPN) dietary treatment. Specific groups are as follows: basal diet as control diet (no yeast), control diet + ADY (2 g/head/d) (ADY2), control diet +ADY (4 g/head/d) (ADY4), and HPN diet (no yeast). The formula design referred to the feeding standard of beef cattle (NY/T 815-2004), in which the control diet (Nemf: 5.11 MJ/kg, CP: 12%) was designed according to the body weight of 150 kg and the daily weight gain of 1.1kg. The HPN diet (Nemf: 5.87 MJ/kg, CP: 13.05%) was designed according to the body weight of 150 kg and the daily weight gain of 1.2 kg.

The beef cattle were given 10 days to adapt to the new environment and the treatment diets before starting the formal experiment. During the adaptation period, the beef cattle were vaccinated following standard operating procedures of the beef facility. The live yeast (2.0×10^10^ CFU/g) was donated by Angel Yeast Co., Ltd, Yichang, Hubei, China. The ingredients and nutritional composition of the basal diets is shown in Table 1.

**Table 1.** Ingredients and nutrient levels of diets (air-dry basis,%)

| Items | Treatments<sup>a</sup> |  |  |  |
| --- | --- | --- | --- | --- |
|  | Control | ADY2 | ADY4 | HPN |
| Corn meal | 26.50 | 26.50 | 26.50 | 39.75 |
| Wheat bran | 3.50 | 3.50 | 3.50 | 5.25 |
| Soybean meal | 2.00 | 2.00 | 2.00 | 3.00 |
| Cottonseed meal | 5.30 | 5.30 | 5.30 | 7.95 |
| Urea | 0.20 | 0.20 | 0.20 | 0.30 |
| Soy oil | 0.20 | 0.20 | 0.20 | 0.30 |
| CaCO<sub>3</sub> | 0.70 | 0.70 | 0.70 | 1.05 |
| CaHPO<sub>4</sub> | 0.20 | 0.20 | 0.20 | 0.30 |
| Na<sub>2</sub>SO<sub>4</sub> | 0.20 | 0.20 | 0.20 | 0.30 |
| NaHCO<sub>3</sub> | 0.40 | 0.40 | 0.40 | 0.60 |
| NaCl | 0.50 | 0.50 | 0.50 | 0.75 |
| Compound premix<sup>b</sup> | 0.30 | 0.30 | 0.30 | 0.45 |
| Distilled grain | 30.00 | 30.00 | 30.00 | 20.00 |
| Hybrid giant napier | 30.00 | 30.00 | 30.00 | 20.00 |
| Total (100%) | 100 | 100 | 100 | 100 |
| Nutritional composition<sup>c</sup> |  |  |  |  |
| NE<sub>mf</sub> /(MJ/kg) | 5.11 | 5.11 | 5.11 | 5.87 |
| CP | 12.00 | 12.00 | 12.00 | 13.05 |
| ADF | 27.55 | 27.55 | 27.55 | 20.46 |
| NDF | 47.77 | 47.77 | 47.77 | 37.57 |
| Ca | 0.73 | 0.73 | 0.73 | 0.78 |
| P | 0.48 | 0.48 | 0.48 | 0.49 |

<sup>a</sup> Treatments: Control, beef cattle fed low plane of nutrition dietary without supplementing active dry yeast; ADY2 and ADY4, beef cattle fed 2 or 4 g/head/d active dry yeast on the control dietary; HPN, beef cattle fed high plane of nutrition dietary without supplementing active dry yeast.
<sup>b</sup> Supplied per kilogram of dietary DM : VA 8000 IU, VD<sub>3</sub> 2000 IU, VE 100 IU, Fe 50 mg, Cu 10 mg, Zn 60 mg, Mn 40 mg, Se 0.2 mg, I 0.5 mg, Co 0.1 mg, Monensin 30 mg.
<sup>c</sup> The value reported for nutritional composition of diets was calculated based on the nutrient analysis from ingredient samples.

### Animals management and growth trial

The growth trial lasted 110 days, and all beef cattle were tethered using neck straps in tie stalls and were individually fed with total mixed ration twice a day (0700 and 1700). During the trial (May to August, 2018), the average ambient temperature was 24.15 ± 4.78°C and the humidity was 64.88 ± 17.67%. The beef cattle were fed *ad libitum* with surplus feeds in trough anytime and had free access to fresh water.

DMI was individually measured based on the differences between the amount of diet offered and refused daily. Body weight was measured before morning feeding on two consecutive days at the beginning and on the final day of the 110-day trial. ADG was calculated as a difference between initial and final live weight and divided by 110 days. Feed conversion ratio (FCR) was calculated as the ratio of individual dietary DMI to ADG.

### Blood sample collection and analysis

Blood samples were collected from the jugular vein of the beef cattle on the morning of the last day of the trial. Tubes were centrifuged at 3200 × g at room temperature for 15 min, the separated serum was stored at −20°C to be used for measurement of UREA, Glucose (GLU), Triglyceride (TG), Total cholesterol (TC), High density lipoprotein (HDL), low density lipoprotein (LDL), non-esterified fatty acid (NEFA), Glutamic-oxaloacetic transaminase (AST), glutamic-pyruvic transaminase (ALT). All serum indexes were determined by a HITACHI-7020 Auto-Biochemical Analyzer.

### Carcass traits and rumen fermentation characteristics

At the end of the experiment, all beef cattle were shipped to a commercial abattoir for slaughter. Hot carcass weight (with kidneys removed), net meat weight, dressing percentage, and net meat percentage were recorded. Dressing percentage was calculated as hot carcass weight (HCW) divided by final BW×100%. Net meat percentage was calculated as net meat weight divided by final BW×100%.

Once slaughtered, the pH of the rumen content was directly measured using a glass electrode pH meter. The rumen fluid was removed from the rumen, filtered with four layers gauze, and placed in a 10 mL centrifuge tube. The collected samples were immediately frozen in liquid nitrogen, and then stored at −80°C until further analysis and the remaining portion was stored at −20°C for volatile fatty acid (VFA), ammonia nitrogen (NH_3_-N), and microbial crude protein (MCP) assay.

### Digestion trial

During the last 4 days (107 d −110 d) of the trial, all feces were collected for 24 h per cattle. The collected feces were weighed and recorded every day. Approximately 100 g of fresh fecal sample from each cattle per day was mixed with 20 mL 10% sulfuric acid and stored at −20°C for further chemical analysis.

### Analytical methods

All chemical analyses were conducted in duplicate. *Longissimus dorsi* muscle samples after being freeze-dried were used to determine Moisture, CP, EE and Ash content. The determination for moisture (method 934.01), CP (method 984.13), EE (method 920.39) and Ash contents (method 942.05) was referenced to standard procedures (AOAC, 2000). The method described by Van Soest et al. [12] was used to determine NDF and ADF, both NDF and ADF were corrected by its ash content. The concentrations of NH_3_-N were measured using phenol-sodium hypochlorite colorimetric[13]. The concentrations of VFAs (acetate, propionate and butyrate) were measured using an HPLC organic acid analysis system (Shimadzu, Kyoto, Japan) The supernatant was shaken with cation exchange resin (Amberlite, IR 120B H AG, ORGANO CORPORATION, Tokyo, Japan) and centrifuged at 6500×g for 5 min. The supernatant was passed through a 0.45 μm filter under pressure, and the filtrate was then injected into an HPLC system. The analytical conditions were as follows: column, SCR-101H (7.9 mm × 30 cm) attached to a guard column SCR(H) (4.0 mm × 5 cm) (Shimadzu); oven temperature, 40°C; mobile phase, 4 mM p-toluenesulfonic acid aqueous solution; reaction phase, 16 mM Bis-Tris aqueous solution containing 4 mM p-toluenesulfonic acid and 100 μM ethylenediaminetetra-acetic acid; flow rate of the mobile and reaction phase, 0.8 mL/1min; detector, conductivity detector (CDD-6A, Shimadzu). The MCP concentration were quantified using a BCA Protein Assay Kit manufactured by Nanjing Jiancheng Bioengineering Institute.

### Bioinformatics analyses (DNA extraction, sequencing)

Total genomic DNA was extracted from the samples using a DNeasy Power Soil Kit (Qiagen, Valencia, CA, USA) according to the manufacturer’s instructions. DNA concentration and quality were checked using a Nano Drop Spectrophotometer. DNA was diluted to 10 ng/μL using sterile ultrapure water and stored at −80°C for downstream use. PCR Primer 16S V4: 515F (5’-GTGCCAGCMGCCGCGGTAA-3’) and 806R(5’-GGACTACHVGGGTWTCTAAT-3’) 16S rRNA genes were amplified using the specific primer with 12 nt unique barcode[14, 15]. The PCR mixture (25 μL) contained 1 × PCR buffer, 1.5 mM MgCl_2_, each deoxynucleoside triphosphate at 0.4 μM, each primer at 1.0 μM, 0.5 U of KOD-Plus-Neo (TOYOBO) and 10 ng template DNA. The PCR amplification program consists of initial denaturation at 94°C for 1 min, followed by 30 cycles (denaturation at 94°C for 20 s, annealing at 54°C for 30 s, and elongation at 72°C for 30 s), and a final extension at 72°C for 5 min. Three replicates of the PCR reactions for each sample were combined and the PCR products were purified using Gel Extraction Kit (Omega Bio-Tek, USA). DNA was quantified using Qubit@ 2.0 Fluorometer (Thermo Scientific). PCR products from different samples were pooled with equal molar amount. Library preparation and sequencing libraries were generated using TruSeq DNA PCR-Free Sample Prep Kit following manufacturer’s recommendations and index codes were added. The library quality was assessed on the Qubit@ 2.0 Fluorometer (Thermo Scientific) and Agilent Bioanalyzer 2100 system. At last, the library was applied to paired-end sequencing (2×250 bp) with the Illumina Hiseq apparatus at Rhonin Biosciences Co., Ltd.

To maintain the Phred quality score of the reads, low-quality sequences were trimmed using Trimmomatic and Usearch before assembly with the paired-end assembler[16, 17]. UPARSE was used to cluster the sequences into OTUs (operational taxonomic units) as well as choose the representative sequence of each OTU at 97% similarity followed by the removal of chimeras and singletons by UCHIME[16]. Four alpha diversity indices (Simpson, Shannon-Wiener, Chao1 and phylogenetic distance) were calculated. Principal component analysis (PCA) was applied to reduce the dimensions of original community data. The OTU table, rarefaction dilution curves, and beta diversity analysis were performed using R programming tools (version 3.3.0).

### Statistical Analysis

The Mixed model (SAS 9.4 Institute Inc., Cary, NC, USA) included treatment as fixed effect and the beef cattle as random effect. For repeated measures, various covariance structures were tested with the final choice exhibiting the lowest value for Akaike’s information criteria. Differences between treatments were declared significant at *P*<0.05.

## Results

### Growth Performance

Effect of dietary supplementation of ADY on growth performance of beef cattle is shown in Table 2. The HPN group had greater final body weight than control (*P*=0.021). The yeast addition had no significant effect on final body weight compared with control (*P*>0.05). However, the ADY group also had no significant difference with HPN (*P*>0.05). Compared to control, supplementation of ADY increased ADG (*P*<0.001), and no significant difference was obtained between ADY4 and HPN group (*P*>0.05). The DMI of HPN group was greater than control and ADY2 group (*P*=0.017), and no significant difference was observed between ADY and control (*P*>0.05), and the DMI of ADY4 group had no significant difference with HPN group (*P*>0.05). No significant effect was detected on FCR among the four groups (*P*=0.262).

**Table 2.** Effect of dietary supplementation of active dry yeast on growth performance of beef cattle

| Items | Treatments<sup>a</sup> |  |  |  | SEM | P-Value |
| --- | --- | --- | --- | --- | --- | --- |
|  | Control | ADY2 | ADY4 | HPN |  |  |
| Initial body weight (kg) | 107.44 | 108.50 | 109.06 | 117.25 | 4.547 | 0.415 |
| Final body weight (kg) | 215.72<sup>b</sup> | 221.87<sup>ab</sup> | 224.56<sup>ab</sup> | 237.29<sup>a</sup> | 4.661 | 0.021 |
| Average daily gain (kg) | 0.98<sup>c</sup> | 1.03<sup>b</sup> | 1.05<sup>ab</sup> | 1.09<sup>a</sup> | 0.011 | <0.001 |
| Dry matter intake (kg) | 5.69<sup>b</sup> | 5.72<sup>b</sup> | 5.97<sup>ab</sup> | 6.95<sup>a</sup> | 0.275 | 0.017 |
| Feed conversion ratio | 5.74 | 5.67 | 5.68 | 6.35 | 0.267 | 0.262 |

Different superscript letters within a row represent significant differences ( $P < 0.05$ ).
<sup>a</sup> Treatments: Control, beef cattle fed low plane of nutrition dietary without supplementing active dry yeast; ADY2 and ADY4, beef cattle fed 2 or 4 g/head/d active dry yeast on the control dietary; HPN, beef cattle fed high plane of nutrition dietary without supplementing active dry yeast.

### Blood indexes

Effect of dietary supplementation of ADY on blood biochemical indices of beef cattle is presented in Table 3. Compared to HPN group, the concentration of UREA in ADY4 group was lowered (*P*=0.005). However, the ADY4 group was not significantly different from the control and ADY2 (*P*>0.05). The concentration of LDL in ADY4 group was higher than HPN group (*P*=0.033), no difference was observed among control and ADY group. The activity of ALT in control and ADY4 groups were higher than HPN group (*P*=0.001), no difference was observed among control and ADY group (*P*>0.05). Supplementation of ADY and HPN diet had no significant effect on concentration of GLU, TG, TC, HDL, NEFA and AST activity (*P*>0.05).

**Table 3.** Effect of dietary supplementation of active dry yeast on blood biochemical indices of beef cattle

| Items<sup>b</sup> | Treatments<sup>a</sup> |  |  |  | SEM | P-value |
| --- | --- | --- | --- | --- | --- | --- |
|  | Control | ADY2 | ADY4 | HPN |  |  |
| UREA (mmol/L) | 5.31<sup>ab</sup> | 5.17<sup>ab</sup> | 5.03<sup>b</sup> | 5.45<sup>a</sup> | 0.079 | 0.005 |
| GLU (mmol/L) | 2.85 | 3.21 | 2.98 | 3.17 | 0.153 | 0.345 |
| TG (mmol/L) | 0.21 | 0.19 | 0.17 | 0.15 | 0.012 | 0.054 |
| TC (mmol/L) | 2.22 | 2.36 | 2.56 | 2.23 | 0.172 | 0.569 |
| HDL (mmol/L) | 1.04 | 1.16 | 1.17 | 0.99 | 0.139 | 0.168 |
| LDL (mmol/L) | 0.20<sup>ab</sup> | 0.21<sup>ab</sup> | 0.26<sup>a</sup> | 0.17<sup>b</sup> | 0.019 | 0.033 |
| NEFA (mmol/L) | 0.05 | 0.05 | 0.05 | 0.04 | 0.006 | 0.715 |
| AST (U/L) | 54.17 | 50.00 | 47.33 | 44.33 | 2.560 | 0.059 |
| ALT (U/L) | 23.00<sup>a</sup> | 20.33<sup>ab</sup> | 24.00<sup>a</sup> | 17.50<sup>b</sup> | 1.052 | 0.001 |

Different superscript letters within a row represent significant differences ( $P<0.05$ ).
<sup>a</sup> Treatments: Control, beef cattle fed low plane of nutrition dietary without supplementing active dry yeast; ADY2 and ADY4, beef cattle fed 2 or 4 g/head/d active dry yeast on the control dietary; HPN, beef cattle fed high plane of nutrition dietary without supplementing active dry yeast.
<sup>b</sup> GLU, glucose; TG, triglyceride; TC, total cholesterol; HDL, high density lipoprotein; LDL, low density lipoprotein; NEFA, non-esterified fatty acid; AST, glutamic-oxalacetic transaminase; ALT, glutamic-pyruvic transaminase.

### Nutrient Apparent Digestibility

Effect of dietary supplementation of ADY on nutrient apparent digestibility of beef cattle is displayed in Table 4. The CP digestibility in HPN group was greater than control and ADY2 group (*P*=0.001), and no difference was obtained between ADY4 and HPN group (*P*>0.05). The ADF (*P*=0.043) and NDF (*P*=0.022) digestibility in HPN group were greater than control group, and no difference was obtained among the ADY2, ADY4 and HPN group (*P*>0.05).

**Table 4.** Effect of dietary supplementation of active dry yeast on nutrient apparent digestibility of beef cattle

| Items | Treatments<sup>a</sup> |  |  |  | SEM | P-value |
| --- | --- | --- | --- | --- | --- | --- |
|  | Control | ADY2 | ADY4 | HPN |  |  |
| DM (%) | 67.50 | 70.23 | 70.16 | 68.92 | 1.144 | 0.314 |
| CP (%) | 52.24<sup>b</sup> | 52.71<sup>b</sup> | 53.48<sup>ab</sup> | 54.45<sup>a</sup> | 0.344 | 0.001 |
| EE (%) | 67.26 | 70.60 | 70.78 | 67.59 | 1.512 | 0.228 |
| ADF (%) | 43.71<sup>b</sup> | 46.55<sup>ab</sup> | 46.85<sup>ab</sup> | 50.59<sup>a</sup> | 1.565 | 0.043 |
| NDF (%) | 46.60<sup>b</sup> | 51.47<sup>ab</sup> | 51.67<sup>ab</sup> | 52.58<sup>a</sup> | 1.344 | 0.022 |

Different superscript letters within a row represent significant differences ( $P<0.05$ ).
<sup>a</sup> Treatments: Control, beef cattle fed low plane of nutrition dietary without supplementing active dry yeast; ADY2 and ADY4, beef cattle fed 2 or 4 g/head/d active dry yeast on the control dietary; HPN, beef cattle fed high plane of nutrition dietary without supplementing active dry yeast.

### Ruminal Fermentation Characteristics

Table 5 shows the effect of dietary supplementation of ADY on ruminal fermentation parameters of beef cattle. The concentrations of NH_3_-N was lowered by ADY supplementation (*P*=0.003) compared with control. The concentration of total VFA (*P*=0.001) and acetic acid (*P*=0.019) in HPN group were greater than ADY4 group. However, no difference was obtained among the control, ADY2 and ADY4 group (*P*>0.05). The concentration of propionic acid in ADY2, ADY4 and HPN group were greater than control (*P*<0.001). No significant difference was observed in pH, MCP and Acetic acid /Propionic acid (*P*>0.05).

**Table 5.** Effect of dietary supplementation of active dry yeast on ruminal fermentation parameters of beef cattle

| Items | Treatments<sup>a</sup> |  |  |  | SEM | P-value |
| --- | --- | --- | --- | --- | --- | --- |
|  | Control | ADY2 | ADY4 | HPN |  |  |
| pH | 5.85 | 6.18 | 6.33 | 6.21 | 0.057 | 0.185 |
| NH<sub>3</sub>-N (mg/100mL) | 7.70<sup>a</sup> | 7.42<sup>b</sup> | 7.33<sup>b</sup> | 7.45<sup>ab</sup> | 0.065 | 0.003 |
| MCP (mg/dL) | 206.07 | 217.32 | 223.36 | 220.85 | 6.121 | 0.282 |
| Total VFA (mmol/L) | 74.82<sup>ab</sup> | 75.13<sup>ab</sup> | 73.12<sup>b</sup> | 76.63<sup>a</sup> | 0.817 | 0.001 |
| Acetic acid (mmol/L) | 53.01<sup>ab</sup> | 52.58<sup>ab</sup> | 49.99<sup>b</sup> | 53.56<sup>a</sup> | 0.775 | 0.019 |
| Propionic acid (mmol/L) | 12.58<sup>b</sup> | 13.67<sup>a</sup> | 14.02<sup>a</sup> | 13.82<sup>a</sup> | 0.180 | <0.001 |
| Butyric acid (mmol/L) | 9.23 | 8.88 | 9.11 | 9.24 | 0.167 | 0.610 |
| Acetic acid/Propionic acid | 4.21 | 3.83 | 3.52 | 3.88 | 0.092 | 0.235 |

Different superscript letters within a row represent significant differences ( $P<0.05$ ).
<sup>a</sup> Treatments: Control, beef cattle fed low plane of nutrition dietary without supplementing active dry yeast; ADY2 and ADY4, beef cattle fed 2 or 4 g/head/d active dry yeast on the control dietary; HPN, beef cattle fed high plane of nutrition dietary without supplementing active dry yeast.

### **S**laughter Performance

As shown in Table 6, the live weight before slaughter of HPN group were greater than control (*P*=0.023), but there was no significant difference observed among ADY2, ADY4 and HPN group. The carcass weight of HPN group was significantly greater than control (*P*=0.001). However, no significant difference obtained between ADY4 and HPN group. The net meat weight of HPN group was significantly greater than the other three groups (*P*<0.001). No significant difference was obtained in dressing percentage (*P*=0.448) and net meat percentage (*P*=0.346) among the four groups.

**Table 6.**
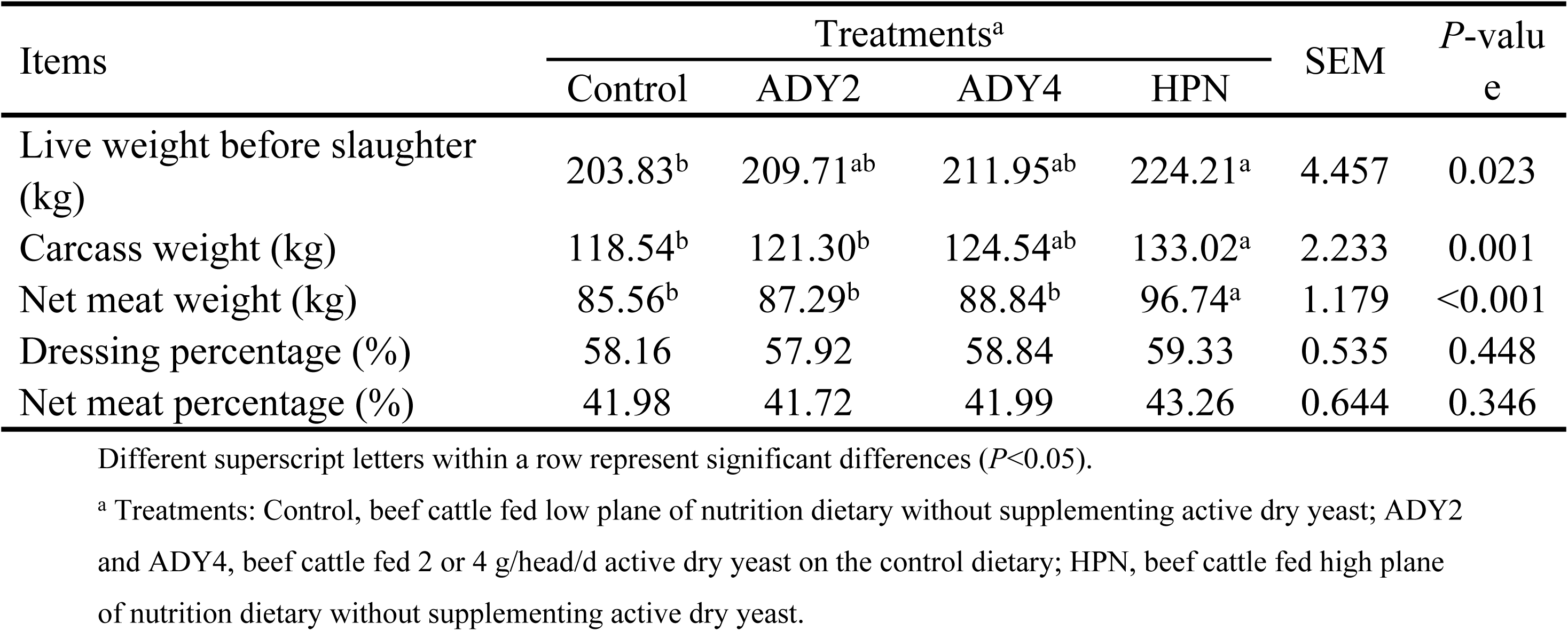
Effect of dietary supplementation of active dry yeast on slaughter performance of beef cattle

### Chemical Composition of Meat

As shown in Table 7, compared with control group, neither ADY supplementation group nor HPN group had significant effects on beef Moisture, CP, EE and Ash content (*P*>0.05).

**Table 7.** Effect of dietary supplementation of active dry yeast on chemical composition in sirloin steak of beef cattle

| Items | Treatments<sup>a</sup> |  |  |  | SEM | P-value |
| --- | --- | --- | --- | --- | --- | --- |
|  | Control | ADY2 | ADY4 | HPN |  |  |
| Moisture (%) | 72.46 | 74.10 | 72.00 | 73.27 | 0.685 | 0.175 |
| CP (%) | 19.47 | 19.99 | 19.70 | 20.32 | 0.519 | 0.685 |
| EE (%) | 2.33 | 2.32 | 2.34 | 2.42 | 0.193 | 0.760 |
| Ash (%) | 3.68 | 4.04 | 3.66 | 4.09 | 0.142 | 0.438 |

<sup>a</sup> Treatments: Control, beef cattle fed low plane of nutrition dietary without supplementing active dry yeast; ADY2 and ADY4, beef cattle fed 2 or 4 g/head/d active dry yeast on the control dietary; HPN, beef cattle fed high plane of nutrition dietary without supplementing active dry yeast.

### Rumen Bacterial Communities

Rarefaction curves obtained from bacterial sequences is shown in Fig 1. As can be seen from Fig 1, the curve of each sample was nearly asymptotic, which indicated that its sequencing depth has substantially covered the entire microbial population. In total, 804875 bacterial sequences were produced, and based on 97% sequence similarity, the sequences were clustered into 32215 OTUs. Distribution of valid sequences and OTUs were presented in supplementary Table S1. The alpha diversity indexes (Chao1, phylogenetic distance, Simpson and Shannon) of the rumen bacterial community in each group are presented in Fig 2. The Chao1 index and the phylogenetic distances index did not differ among the four treatments, but the Simpson (*P*=0.014) and Shannon (*P*=0.045) indices of the control and HPN group were greater than ADY4 group. It can be seen from Fig 3 that the first and second principal components based on principal component analysis (PCA) respectively explain the bacterial community structure variation of 18.70% and 9.60% (Fig 3a), and the first and second principal component based on UniFrac unweighted principal component analysis explained 18.70% and 6.60% of the bacterial community structure variation, respectively(Fig 3b). Based on UniFrac weighted principal component analysis, the first and the second principal component explained 35.4% and 21.3%, respectively (Fig 3c). However, the four treatments did not appear to be independently distributed, and the internal variation between the four groups was large and the similarity was high. At the genus level, ADY4 and HPN group can be clustered together with a distance of 0.04, and minged with ADY2 with a distance of 0.08, then minged with the control with a distance of 0.16, indicating that the microbial composition of ADY4 and HPN have certain similarities (Fig 4).

**Fig 1.**
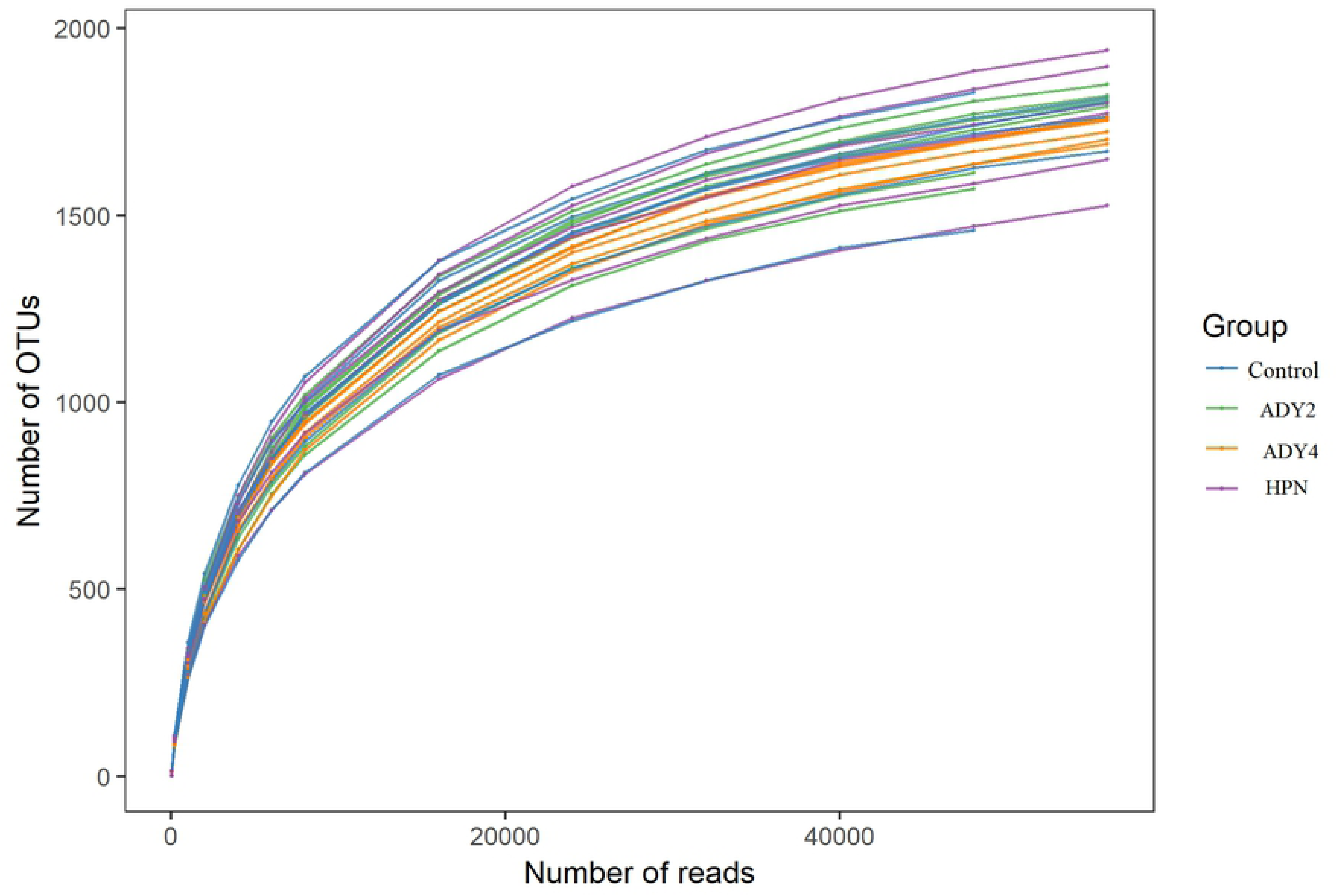
Rarefaction Curve. Rarefaction curves of rumen bacterial communities based on the 16S rRNA gene sequences from the different rumen fluid samples examined at a 0.03 distance level (Control, beef cattle fed low plane of nutrition dietary without supplementing active dry yeast; ADY2 and ADY4, beef cattle fed 2 or 4 g/head/d active dry yeast on the basis of control dietary; HPN, beef cattle fed high plane of nutrition dietary without supplementing active dry yeast).

**Fig 2.**
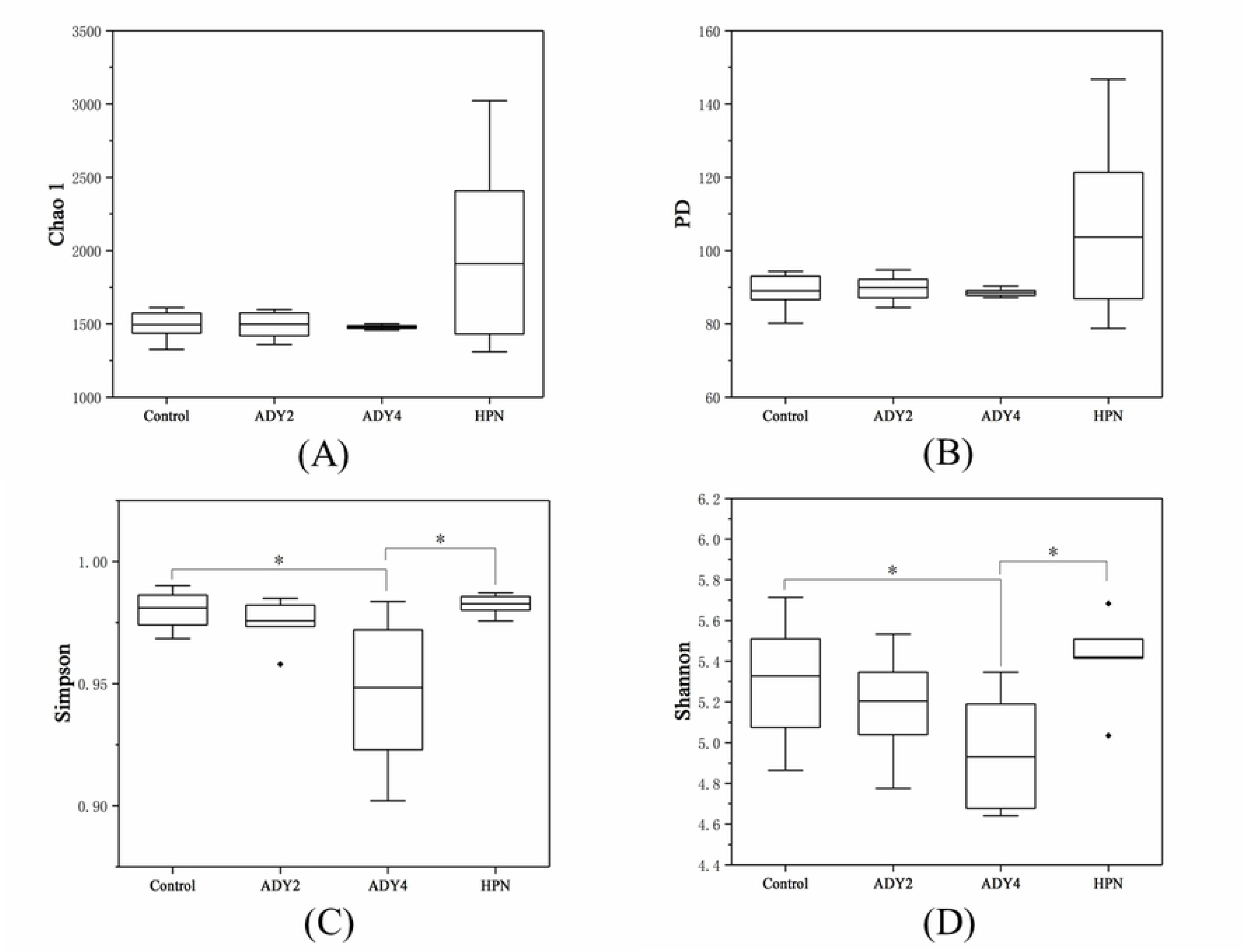
Bacterial alpha diversity indices. (A) Chao1, (B) phylogenetic distance, (C) Simpson, and (D) Shannon index values of the rumen bacterial communities of beef cattle. The samples were labelled Control, ADY2, ADY4, HPN according to different treatments. There are six replicates in each group (Control, beef cattle fed low plane of nutrition dietary without supplementing active dry yeast; ADY2 and ADY4, beef cattle fed 2 or 4 g/head/d active dry yeast on the basis of control dietary; HPN, beef cattle fed high plane of nutrition dietary without supplementing active dry yeast).

**Fig 3.**
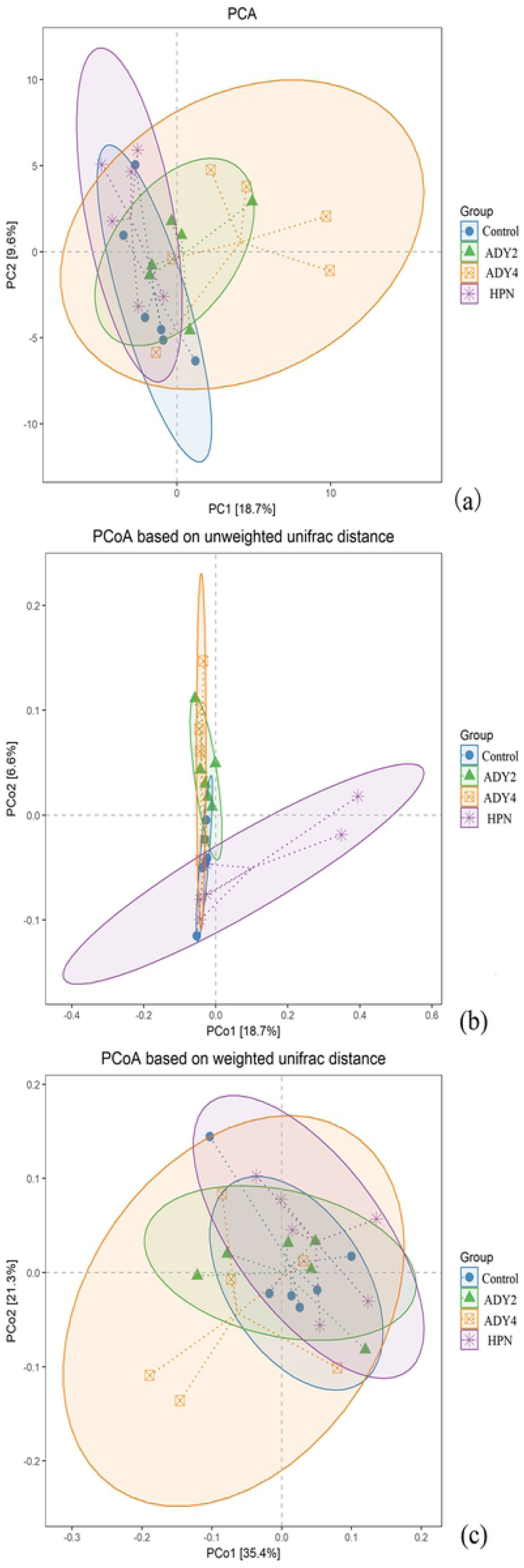
PCA(a), PCoA based on unweighted (b) and weighted (c) unifrac distance shows difference of bacteria community structures among the different treatment groups in beef cattle. Control, beef cattle fed low plane of nutrition dietary without supplementing active dry yeast; ADY2 and ADY4, beef cattle fed 2 or 4 g/head/d active dry yeast on the basis of control dietary; HPN, beef cattle fed high plane of nutrition dietary without supplementing active dry yeast.

**Fig 4.**
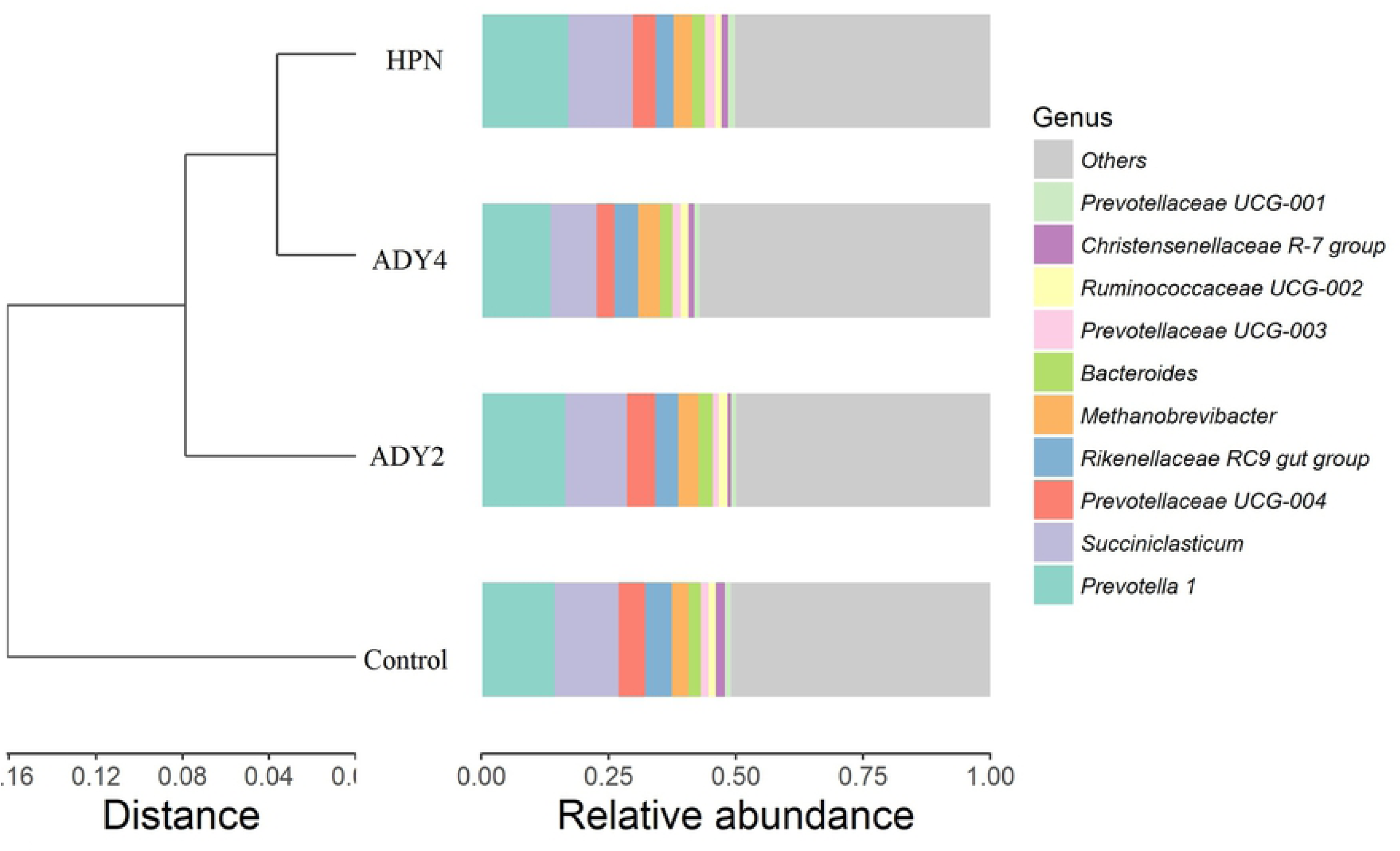
Effect of dietary supplementation of active dry yeast on the genus level of ruminal communities of beef cattle (Control, beef cattle fed low plane of nutrition dietary without supplementing active dry yeast; ADY2 and ADY4, beef cattle fed 2 or 4 g/head/d active dry yeast on the basis of control dietary; HPN, beef cattle fed high plane of nutrition dietary without supplementing active dry yeast).

The microorganism with relatively high abundance in rumen fluid is shown in Table 8. At the phylum level, the relative abundance of Bacteroidetes and Firmicutes was dominating, account for about 85% of the total microorganisms. Compared with the HPN, the relative abundance of Firmicutes in ADY4 group was lowered (*P=*0.015). At the genus level, the relative abundance of *Ruminococcaceae UCG-002* of ADY4 was higher than control and HPN groups (*P=*0.004). In addition, the relative abundance of *Ruminococcaceae UCG-002* in ADY2 group was also higher than the control (*P=*0.004).

**Table 8.**
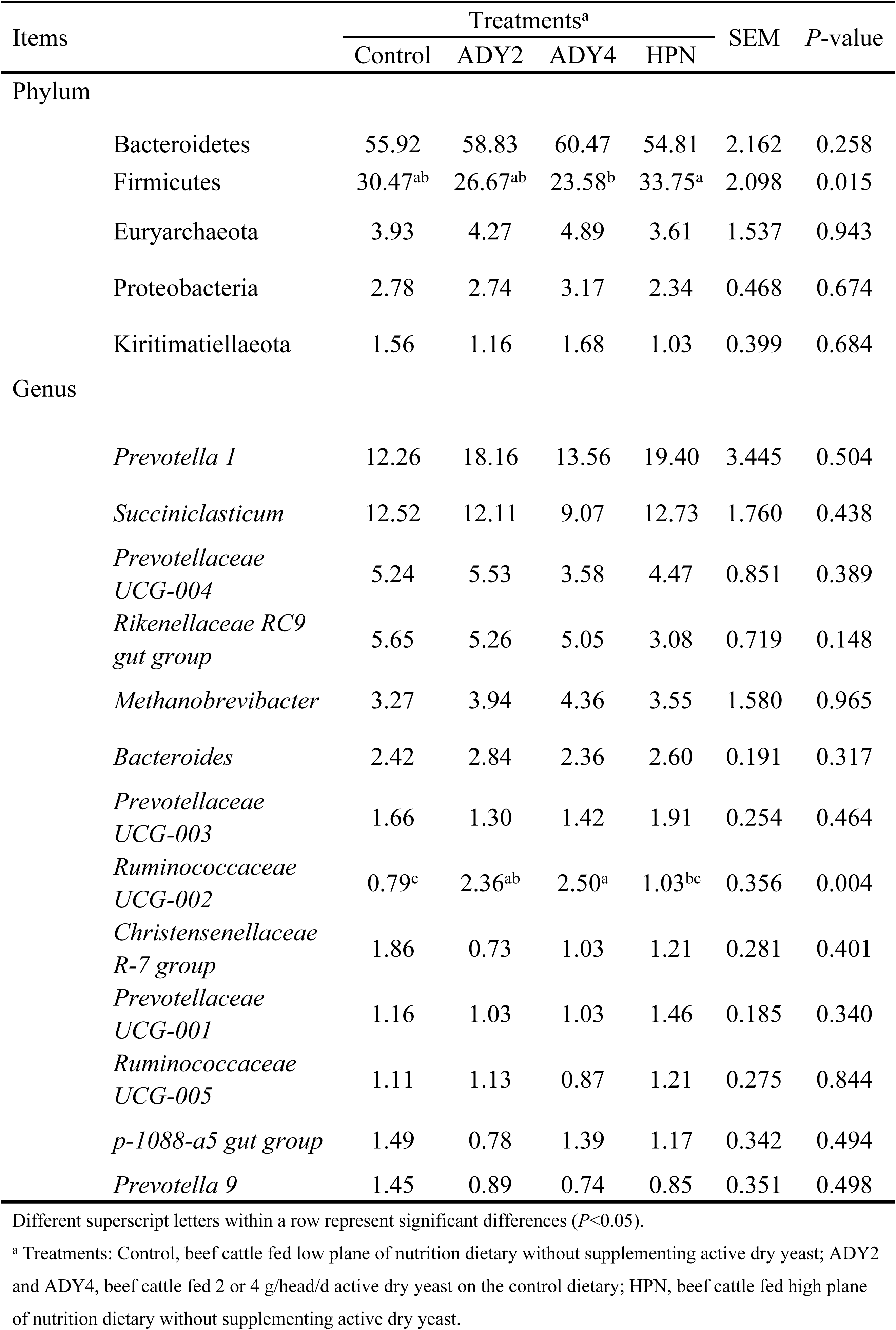
Microorganism with relatively high abundance and significant difference in rumen fluid (%)

## Discussion

As a kind of yeast preparations, ADY is widely used in ruminant animals. Most studies have shown that the addition of yeast preparation to beef cattle diets could improve growth performance. In present study, cattle fed low plane of nutrition diet supplemented with ADY 4 g/head/d significantly increased ADG, and achieved to a comparable level with cattle fed with high plane of nutrition diet, which was in agreement with the research of Geng et al. [18]. Previous studies have shown that the supplementation with ADY could increase ruminant animals DMI[19], however there were also some studies showed that DMI could be reduced[20, 21]. Our results indicated that dietary supplementation with ADY had no significant effect on DMI, which was consistent with the result reported by Vyas et al. [22]. The reason for the inconsistences may be due to the different yeast strains used in the trials[23]. However, the present study found that the DMI of the HPN group was significantly greater than control group. Yuangklang et al. [24] reported that the DMI of beef cattle increased significantly due to the increase of dietary protein. Arriola et al. [25] also reported that increase the amount of concentrate in diets could increase the feed intake of the cattle, which was consistent with our study.

So far, there are few studies focused on the effect of ADY on slaughter performance of beef cattle. The BW of HPN group was significantly higher than the other three low plane nutrition groups no matter ADY supplementation or not. Gleghorn et al. [26] reported that with the increase of CP concentrations in the diet, the carcass weight, dressing percentage and net meat production increased. Our study showed that the addition of ADY increased the final body weight so that its carcass weight increased to a comparable level with the HPN group accordingly, and no significant difference between ADY4 and HPN group was detected, which was consistent with the previous studies[27, 28].

There are many reports focused on the effect of yeast preparations on milk quality, ADY has a certain effect on the milk fat and protein content[29, 30], which indicating that ADY may affect the rumen microbial population. The changes in microbial composition would further alter ruminal VFA production and constitution, which is the reason for the variations of milk fat and protein. In present study, however, no significant effect was detected on the Moisture, CP, EE and Ash content of the *longissimus dorsi* muscle, which was consistent with the study of Geng et al. [18], though the rumen microbial population was also affected by ADY supplementation.

The diagnostic results of serum biochemical analysis provided information on the function and nutritional status of almost all organs, as well as disease progress, which are closely related to the performance of animals. In present study, all serum indexes were not significantly influenced by ADY supplementation, however, the serum UREA concentration in HPN group was significantly greater than ADY4 group, which might indicate that the HPN group cattle could not utilize nitrogen as quickly as ADY4, for the ADY4 group fed low plane of nutrition diet. Findings reported by Geng et al. [18]. showed that the UREA content of finishing bulls could not be affected by ADY supplementation, which was consistent with present study. ALT and AST are important enzymes of liver function, and their activity has a great relationship with the growth and development performance of animals. If the activity of two enzymes in the blood were elevated, it may be caused by liver damage. In this study, compared with the control group, the supplementation of ADY had no significant effects on serum AST and ALT, indicating that ADY had no adverse effect on the liver function. Different from this, reports showed that the addition of ADY could result in fewer liver abscesses[10, 11].

Our results showed that the CP, NDF and ADF digestibility in HPN group was significantly greater than control group, however the yeast addition had no significant effect on CP, NDF and ADF digestibility, though the NDF and ADF digestibility were increased numerically but not statistically. Vyas et al. [31] reported that the addition of probiotics (*Propionibacterium*) to beef cattle diet had no significant effect on the digestibility of DM, NDF and ADF. Sanchez et al. [32] also reported that the addition of probiotics (*Proionibacterium acidipropionici* P1691) to the low-quality coarse fodder had no significant effect on the digestibility of NDF in beef cattle. These results were consistent with our study. It is suggested that adding ADY to the diet could change the composition of rumen bacteria, thereby increasing the digestibility of fiber[33], however the digestibility of NDF and ADF in our study were not increased, probably due to the measurement of total tract apparent digestibility of nutrients, while Jiao et al. reported that the addition of ADY could increased the nutrients digesitiblity in small intestine[9].

NH_3_-N is the main precursor for microbial proteins synthesis in the rumen. The addition of ADY in present study decreased NH_3_-N concentration. This means that the microbe utilize NH_3_-N to synthesize MCP more efficiently. However, Vyas et al. [22] reported that the addition of ADY (4 g/head/d) in the diet increased the concentration of NH_3_-N, probably due to the fluctuating rumen environment leading to the discrepancies. VFA in the rumen, acting as an important source of energy for ruminants, is mainly derived from the fermentation of carbohydrates, which also provides energy for the synthesis of microorganisms in the rumen. Previous study have shown that ADY could not affect the total VFA production in the rumen[34], whereas other study have shown that yeast preparation could increase the proportion of propionic acid, while reduced the proportion of acetic[35] or increased the proportion of acetic acid[36]. In present study, there was no difference in the concentration of rumen total VFA after ADY supplementation, indicating that changes in microbial composition and fermentation capability are too subtle to cause changes in total VFA concentration. However, the VFA concentration in the HPN group was significantly higher than that of the ADY4 group, indicating that there was a gap in the amount of VFA production between the HPN dietary group and the yeast supplementation group. Interestingly, the addition of ADY increased the propionic acid molar proportion, this means that the cattle might utilize energy more efficiently because cattle mainly produces glucose by gluconeogenesis using propionic acid as substtate.

In present study, ADY supplementation reduced the shannon and simpson indices. This was also confirmed by Ogunade et al. [37] in beef cattle. The reduction of shannon and simpson indices was might driven by a reduction in richness coupled with an increase in dominance of some species such as members of Bacteroidetes[38]. At the genus level, *Prevotella* is the dominanting flora in the beef cattle rumen in the four treatment groups, and followed by *Succiniclasticum* and *Prevotellaceae UCG-004* which was consistent with Myer et al. [39]. In present study, the dietary yeast supplementation increased the relative abundance of *Ruminococcaceae UCG-002*, which was consistent with previous studies claiming *Saccharomyces cerevisiae* favor the establishment of cellulolytic bacteria in the rumen[37, 40, 41]. The *Ruminococcaceae* plays key roles in cleaving the cellulose and hemicellulose components of plant material[42], and the increased bacterial number resulting in improved fibre digestibility has been one of the beneficial effects of yeast supplementation in ruminants[43]. Patra et al. [44] reported that the reduction in the relative abundance of *Ruminococcace* would lead to a decrease in fiber digestibility. We also observed increased NDF and ADF digesitiblity at least more than 7% with ADY supplementation, though it was not statistically significant. The increased fibrous material digestibility may attributed to the increased relative abundance of *Ruminococcaceae UCG-002.* At the genus level, clustering analysis indicated that the ADY4 group and HPN group was more similar, suggesting that the addition of ADY shifted the rumen microflora composition fed with low plane of nutrition diet to a high plane of nutrition fed one. Hernandez-Sanabria et al. [45] argured that particular bacteria and their metabolism in the rumen may contribute to differences in host feed efficiency. This was further confirmed by Myer et al. [39] that the growth performance was correlated with the composition of rumen microbiota. The shift of rumen microbial composition partially explained the increased nutrients digestibility and ADG with the ADY supplementation.

## Conclusions

Supplementation ADY 4 g/head/d shifted the microbial compostion of beef cattle fed low plane of nutrition to a more similar level with cattle fed with HPN diet, reduced the Simpson and Shannon indices of beef cattle rumen bacteria, increased the relative abundance of *Ruminococcaceae UCG-002*, and increased the yield of rumen propionate. and increased the ADG of beef cattle. Supplementation ADY 4 g/head/d in low plane of nutrition dietary produced comparable carcass weight to HPN dietary, without adverse effect on the chemical composition of beef.

## Acknowledgements

Author Quanhui Peng thanks for the financial support from 2017YFD0502005, Angel Yeast Co., Ltd, Yichang, Hubei, China, and and Overseas Expertise Introduction Center for Discipline Innovation (“111” center) of Sichuan Agricultural University.

## Supporting information

S1Table. Distribution of valid sequences and OTU per sample.

